# DNAH14 deficiency impairs sperm motility by reducing flagellar beat amplitude

**DOI:** 10.1101/2025.07.28.667185

**Authors:** Yawen Liu, Yanan Zhao, Yali Qiu, Jiaxin Zeng, Haifeng Xu, Bingbing Wu, Xiang Tang, Liying Wang, Wei Li, Chao Liu

## Abstract

Male infertility affects approximately 15-20% of couples worldwide, with asthenozoospermia accounting for 19% of cases. Nevertheless, the genetic basis of many asthenozoospermia cases remains poorly understood. In this study, we generated *Dnah14* knockout mice, which displayed subfertility due to impaired sperm motility despite normal sperm morphology. Comprehensive kinematic analysis revealed that *Dnah14*-deficient spermatozoa exhibited reduced flagellar beat amplitude but increased beat frequency, while other dynein components remained unaffected. Notably, increasing sperm concentration during in vitro fertilization (IVF) experiments partially rescued the fertilization defects caused by *Dnah14* deletion, suggesting a compensatory mechanism for this specific form of asthenozoospermia. Our results establish DNAH14 as a critical regulator of sperm motility and propose a potential therapeutic strategy for affected individuals.

## Introduction

Infertility represents a significant global health challenge, affecting approximately 15-20% of couples worldwide and imposing profound psychosocial and economic burdens^1–3^. Male factors account for 40-50% of infertility cases^2^, with asthenozoospermia accounting for nearly 19% of male infertility diagnoses^4^. According to WHO criteria, asthenozoospermia is defined as sperm with less than 32% progressive motility or 40% total motility^5^. This condition impairs sperm propulsion, severely compromising the ability to navigate the female reproductive tract and fertilize oocytes^6^. The rising prevalence of asthenozoospermia correlates with deteriorating environmental factors, lifestyle changes, genetic factors, and increasing psychological stressors^7^, whereas the causes of many asthenozoospermia cases remains unexplained.

Successful fertilization critically depends on two interdependent sperm attributes: structural integrity and functional motility^8^. Structural integrity ensures proper acrosomal reaction and genetic material delivery, whereas functional motility enables directional navigation through the complex cervico-uterine environment^9^. Flagellar motility is driven by the highly organized “9+2” microtubule axoneme^10^, within which dynein axonemal heavy chain (DNAH) proteins serve as the core molecular motors. These proteins generate sliding forces between microtubule doublets via ATP hydrolysis^11;12^. To date, thirteen DNAH family members have been identified, with mutations in *DNAH1*^12^*, DNAH2*^13^, *DNAH3*^14^, *DNAH5*^15^, *DNAH6*^16;17^, *DNAH8*^18^, *DNAH10*^16^, *DNAH12*^19^, and *DNAH17*^20^ recognized as genetic causes of primary ciliary dyskinesia (PCD) and multiple morphological abnormalities of the flagellum (MMAF) associated asthenoteratozoospermia^21;22^.

DNAH14 is structurally classified as a single-headed inner-arm dynein^23;24^, which is a core mechanochemical component of ciliary and flagellar motility. Biallelic mutations in *DNAH14* were identified in children with PCD, manifesting as chronic respiratory infections, situs inversus, and impaired mucociliary clearance due to defective ciliary beating^25^. Subsequent studies linked DNAH14 dysregulation to cystic fibrosis comorbidities, where it exacerbates pulmonary dysfunction by disrupting airway cilia-mediated pathogen clearance and altering ion transport pathways^26^. Despite these advances in respiratory pathophysiology, the role of DNAH14 in male reproduction remains largely unknown.

In this study, we established a *Dnah14* knockout mouse model to elucidate its role in male reproductive function. Our findings demonstrate that *Dnah14* deletion results in male subfertility primarily through impairment of sperm motility. Comprehensive kinematic analysis revealed that DNAH14 deficiency specifically disrupts sperm flagellar dynamics, characterized by significantly reduced beat amplitude coupled with increased beat frequency. Importantly, immunofluorescence analysis confirmed that the loss of DNAH14 does not compromise the structural integrity or localization of other dynein arm component, indicating a specific rather than global effect on axonemal dynein function. Notably, *in vitro* fertilization (IVF) assays showed that increasing sperm concentrations can partially rescue the fertilization defects in *Dnah14^-/-^* male mice, suggesting that quantitative enhancement may compensate for qualitative motility impairments. These results provide mechanistic insights into the pathogenesis of asthenozoospermia and identify DNAH14 as a critical regulator of sperm movement.

## Materials and Methods

### Animals

The *Dnah14* knockout mouse model was generated by CRISPR/Cas technology using the gRNA target sequence (gRNA1: GAGGTTCTTACTGTAACAAAAGG; gRNA2: CCCTTGCAGCCTAGTTGTCAAGG). We obtained *Dnah14* knockout mice through the deletion of exon 5 to exon 10 of the *Dnah14* gene. The mice were genotyped by PCR and DNA sequencing analysis. The PCR primers used for genotyping were as follows: F1, 5’-TACTCTGCTCTTCTGCGTTCATA-3’; R1, 5’-GACTACCCTTTCTTTCATAGTCTTCAC-3’; R2, 5’-CCCTACACATTGTGGACTGAACTTT-3’. The wild-type allele generated a 626 bp band, the homozygotes produced a 466 bp band, and the heterozygotes produced 626 bp and 466 bp bands. All animal experiments and experimental protocols were conducted in accordance with the guidelines established by the institutional animal care and use committee (IACUC). The IACUC protocols (Ethics No. RSDW-2024-01391) of the Laboratory Animal Welfare and Ethics Committee of Guangzhou Women and Children’s Medical Center.

### Phylogenetic analysis

Amino acid sequences of DNAH14 of 13 species were downloaded from NCBI (https://www.ncbi.nlm.nih.gov/protein) and UniProt (http://www.uniprot.org). The phylogenetic trees were constructed using MEGA 11^27^ with the Neighbor-Joining method^28^. Bootstrap analyses were carried out using 1000 replications with the p-distance model^29^.

### RNA extraction and RT-PCR

Total RNA was extracted from tissues using TRIzol reagent. cDNA was synthesized by the PrimeScript^TM^ RT Reagent Kit (RR037A, TaKaRa, Beijing, China). The expression level of *Dnah14* in the tissue was analyzed by RT-PCR using the primers (F: 5’-AAAAGAGACACTCGCTGCTGA-3’ and R: 5’-TTGAGTGTTTCCTCGGCCTTTC-3’).

Here, *gapdh* was used as a control gene using the primers (F: 5’-TGCATTTGGACCATTGGTC-3’ and R: 5’-TTTGCACTGGTAACGTGTTCAT-3’).

RT-PCR was performed under the following conditions: 94°C for 4 min, 25 cycles at 94°C for 30 s, 58°C for 30 s and 72°C for 20 s, and 72°C for 10 min.

### Assessment of fertility

Fertility was assessed in 2-month-old male mice of various genotypes (four mice per genotype). Each male was paired with two wild-type females (6-8 weeks of age) for a period of 6 months. The time of parturition and litter size were recorded, and the number of litters was tallied every two weeks.

### Tissue phenotype and histological analysis

After recording body weight, male mice were euthanized by cervical dislocation. Tissue samples designated for mRNA and protein analysis were collected and immediately flash-frozen in liquid nitrogen. Testes and cauda epididymides were fixed overnight in Bouin’s solution or 4% paraformaldehyde (PFA), respectively. Fixed tissues were subsequently dehydrated, cleared, and embedded in paraffin. Sections (5 μm) of paraffin-embedded testes and cauda epididymides were stained with H&E and PAS-haematoxylin using established protocols^30^.

### Epididymal sperm count

The unilateral cauda epididymis was dissected from mice, finely minced in 1 ml of pre-warmed human tubal fluid (HTF) medium (Millipore), and incubated for 10 min at 37°C in a 5% COL atmosphere. Spermatozoa released into the medium were quantified by counting three separate replicates using a hemocytometer on a Primo Star microscope (ZEISS).

### Immunofluorescence

Spermatozoa were released from the cauda epididymides of 2-month-old *Dnah14^+/+^* and *Dnah14^-/-^* mice by incubation in PBS at 37°C for 10 min. The collected spermatozoa were then spread onto glass slides for morphological assessment or immunostaining. After air-drying, samples were fixed with 4% PFA for 5 min at room temperature (RT), followed by three PBS washes. Slides were blocked in 5% bovine serum albumin (BSA) for 30 min at RT. Primary antibodies were applied to the sperm preparations and incubated overnight at 4°C. After washing, secondary antibodies were applied and incubated. Nuclei were counterstained with DAPI, and images were acquired using an ECLIPSE Ti2 microscope (Nikon).

### Antibodies

The primary antibodies used for immunoblotting or immunofluorescence analysis of various proteins in this study were as follows: Mouse anti-β-TUBULIN (1: 400, M20005L, Abmart), rabbit anti-β-TUBULIN (1: 400, A12289, Abclonal), mouse anti-SP56 (1: 200, 55101, QED), rabbit anti-AKAP4 (1: 200, A14813, Abclonal), rabbit anti-septin4 (SEPT4, 1: 200, A10238, Abclonal), rabbit anti-DNAH6 (1: 100, 18080-1-ap, Proteintech), rabbit anti-DNAH17 (1: 100, 24488-1-ap, proteintech) were used for immunofluorescence. The secondary antibodies were goat anti-rabbit FITC (1: 200, ZF-0311, Zhong Shan Jin Qiao) and goat anti-mouse TRITC (1: 200, ZF-0313, Zhong Shan Jin Qiao) for immunofluorescence. Additionally, the following reagents were used for immunofluorescence: PK Mito Deep Red (1: 2000 dilution, PKMDR-1, Genvivo).

### Transmission electron microscopy

Adult testis tissues were fixed overnight in a solution containing 1.5% glutaraldehyde, 1.5% paraformaldehyde (PFA), and 0.1 M cacodylic acid sodium salt trihydrate. Fixed samples were then trimmed, washed with 0.1 M cacodylate buffer, and post-fixed in 1% osmium tetroxide (OsO_4_). Tissues were dehydrated through a graded acetone series, infiltrated with Epon resin mixtures, and embedded in pure resin. Ultrathin sections were imaged using a transmission electron microscope (JEM-1400, JEOL, Tokyo, Japan).

### Sperm motility

Spermatozoa were released from incised cauda epididymides into phosphate-buffered saline (PBS, Gibco, C14190500BT) and incubated for 10 min at 37°C. The resulting swim-up suspension was assessed for sperm motility using an Olympus BX51 microscope (Japan) with a 20× phase-contrast objective. Fields of view were recorded via a CCD camera (Olympus), followed by computer-assisted semen analysis (CASA) using the Minitube Sperm Vision Digital Semen Evaluation System (Model 12500/1300; Minitube Group, Tiefenbach, Germany).

### Flagellar movement of tethered sperm head

Flagellar waveform analysis was performed via head tethering for planar beating as previously described. Briefly, spermatozoa from dissected cauda epididymides of adult *Dnah14^+/+^* and *Dnah14^-/-^*males were incubated for 10 min in pre-warmed HTF medium at 37°C. Subsequently, 10 µl of sperm suspension was placed on a slide and gently covered to immobilize sperm heads. Flagellar movements of tethered spermatozoa were recorded at 37°C for 1 s at 60 frames per second (fps) using an inverted microscope equipped with a high-speed camera. Dark-field imaging was conducted at 200 fps with a collimated LED light source. For analysis, images underwent Gaussian blur processing (σ: 0.5 pixels) and background subtraction (radius: 5 pixels) in ImageJ. Flagellar dynamics were quantified using SpermQ^31^, defining positive curvature as bending toward the hook-shaped head’s principal bend direction. Peak frequency was derived from Fast Fourier Transform (FFT) analysis of curvature values.

### *In vitro* fertilization

4-week-old wild-type female mice were intraperitoneally injected with 5 IU of pregnant mare serum gonadotropin (PMSG, #110914564, NINGBO SANSHENG, China). After 46-48 hours, the mice received an intraperitoneal injection of 5 IU human chorionic gonadotropin (hCG, #110911282, NINGBO SANSHENG, China). 15 hours post-hCG administration, oocyte-cumulus complexes (OCCs) were surgically collected from the ampulla of the oviducts. Spermatozoa were isolated from the cauda epididymis of 9-week-old *Dnah14*^+/+^ and *Dnah14^-/-^* male mice and released into 200 μL of human tubal fluid (HTF) medium (Sudgen Biotechnology, China). Sperm capacitation was performed by incubation at 37°C in a 5% CO_2_ incubator for 1 h. Meanwhile, cumulus-free oocytes were cultured in 100 μL HTF medium under mineral oil (M5310, Sigma-Aldrich) at 37°C in a 5% CO_2_ atmosphere for in vitro fertilization. For IVF, capacitated sperm (1-4 × 10^5/mL) were co-incubated with ovulated oocytes in HTF medium for 3 h. Fertilized oocytes were then washed in fresh HTF droplets to remove abnormal oocytes, with all procedures conducted on a temperature-controlled plate (TPiE-SMZR, TOKAI HIT) maintained at 37°C. Successful fertilization was confirmed by the presence of male and female pronuclei 6 h post-insemination, and the 2-cell embryo development rate was assessed at 24 h post-fertilization.

### Statistical analysis

All statistical analyses were conducted using GraphPad Prism software (version 8.02, San Diego, USA) utilizing unpaired two-tailed Student’s t tests. Error bars represent the standard error of the means (SD) with a ± sign. Statistical significance was determined when the *P* value was less than 0.05 (*), 0.01 (**) or 0.001 (***).

## Results

### DNAH14 deficiency leads to male subfertility

Phylogenetic analysis revealed strong conservation of the *Dnah14* gene across mammalian species, with notable divergence in avian lineages (Fig. 1A). To investigate the physiological role of DNAH14 in male reproduction, we first characterized *Dnah14* expression pattern in mouse different tissues and found it was predominantly expressed in testis (Fig. 1B). Further analysis of mouse testes from different days after birth was carried out. *Dnah14* was highly expressed from postnatal day 28 onward (Fig. 1C), supporting the potential functional role of DNAH14 during spermiogenesis. Using CRISPR/Cas9-mediated genome editing, we generated a constitutive *Dnah14* knockout mouse line by deleting a 16,293-bp genomic region spanning exon 5 to 10 (Fig. 1D). *Dnah14^-/-^* mice were genotyped (Fig. 1E) and confirmed by polymerase chain reaction (PCR), further qRT-PCR analysis verified the absence of *Dnah14* transcripts in *Dnah14^-/-^* testes (Fig. 1F). *Dnah14^-/-^* males exhibited normal somatic development, with no differences in body weight or gross morphology compared to *Dnah14^+/+^* male mice (Fig. 1J). However, fertility testing showed that average litter sizes were significantly decreased in *Dnah14^-/-^*male mice compared to control males (Fig. 1G), indicating the DNAH14 deficiency causes male subfertility.

**Figure 1.**
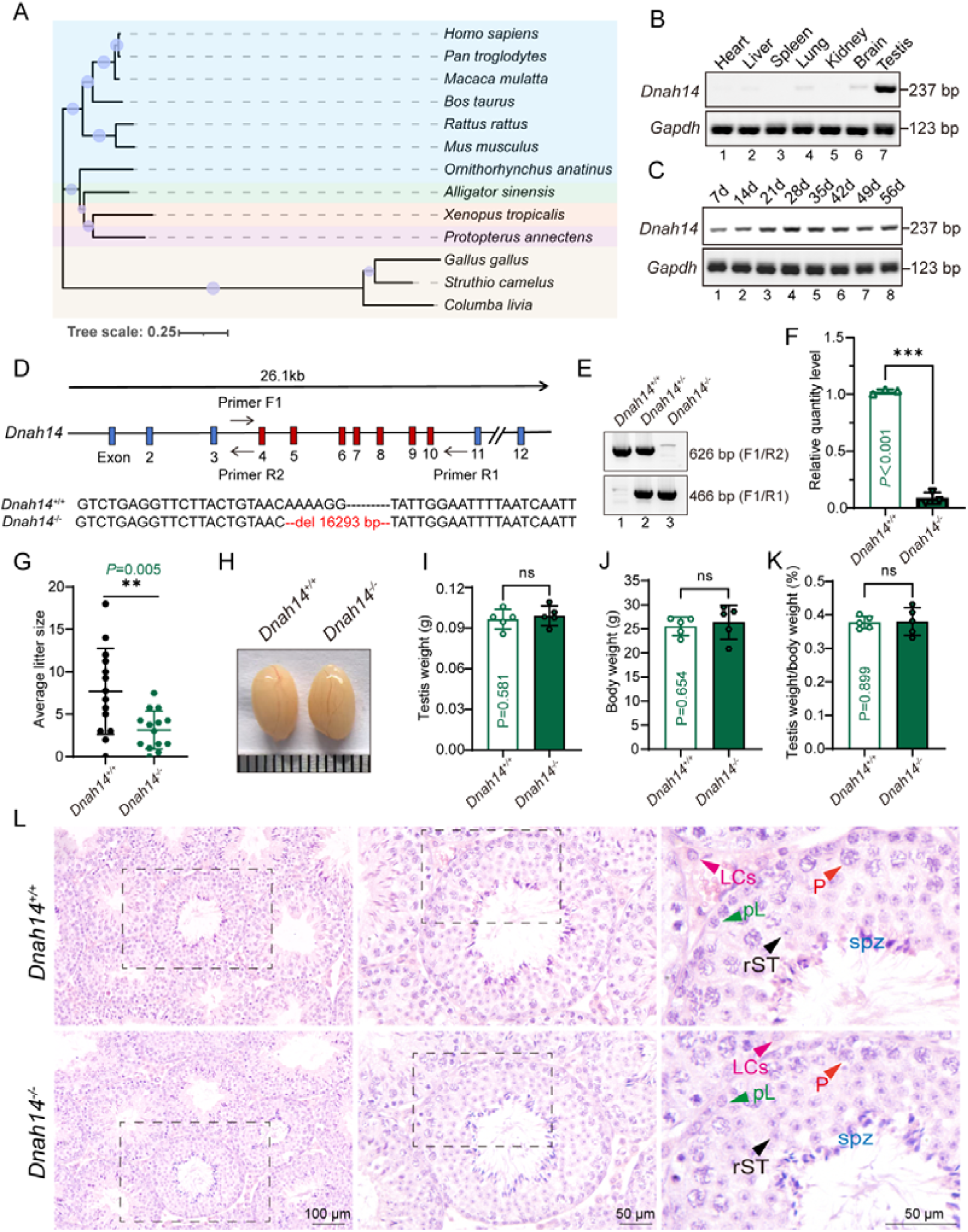
*Dnah14* knockout reduces male feritility in mice. (A) Multiple species phylogenetic tree of DNAH14. Structural similarity scores (Z scores) of DNAH14 orthologs in *Homo. sapiens, Pan troglodytes, Macaca mulatta, Bos taurus, Rattus rattus, Mus musculus, Ornithorhynchus anatinus, Alligator sinensis, Xenopus tropicalis, Protopterus annectens, Gallus gallus, Struthio camelus* and *Columba livia*. (B) RNA expression profile of *Dnah14* in various mouse tissues, with *Gapdh* as a loading control. (C) RNA expression pattern of *Dnah14* in mouse testes at different postnatal days, with *Gapdh* as a loading control. (D) Schematic diagram of *Dnah14^-/-^* mouse construction via CRISPR-Cas9 mediated genome editing. (E) Genotyping to identify *Dnah14* knockout mice. (F) Quantification of *Dnah14* mRNA levels in testes of *Dnah14^+/+^*and *Dnah14^-/-^* mice (n=3 independent experiments). Data are presented as the means ± SD. Two-tailed Student’s t test, \*\*\**P*<0.001. (G) Reduced fertility in *Dnah14* knockout male mice. Four groups of *Dnah14^+/+^* and *Dnah14^-/-^*mice were each paired with two wide type female mice, and litter sizes were recorded over six months. Data are presented as the means ± SD. Two-tailed Student’s t test, \*\**P*=0.005. (H) Representative images of testes from 2-month-old *Dnah14^+/+^* and *Dnah14^-/-^* mice. (I) Quantification of testis weight in *Dnah14^+/+^* and *Dnah14^-/-^* male mice (n=5 independent experiments). Data are presented as the means ± SD. Two-tailed Student’ s t test; ns: no significance. (J) Quantification of body weight in *Dnah14^+/+^*and *Dnah14^-/-^* male mice (n=5 independent experiments). Data are presented as the means ± SD. Two-tailed Student’ s t test; ns: no significance. (K) Ratio of testis weight to body weight (%) in *Dnah14^+/+^*and *Dnah14^-/-^* male mice (n=5 independent experiments). Data are presented as the means ± SD. Two-tailed Student’ s t test; ns: no significance. (L) Hematoxylin-eosin (H&E) staining of testis sections from *Dnah14^+/+^* and *Dnah14^-/-^* mice. LCs: Leydig cells, pL: Preleptotene spermatocyte, P: Pachytene spermatocyte, rSt: round spermatid, eSt: elongating spermatid. Scale bars=100 μm, 50 μm.

### *Dnah14* knockout mice show normal spermatogenesis

To investigate the cause of male subfertility, we examined *Dnah14^-/-^*testis at both macroscopic and histological levels. *Dnah14* knockout did not affect testis size (Fig. 1H), testis weight (Fig. 1I), and the ratio of testis weight to body weight (Fig. 1K). Hematoxylin and eosin (H&E) staining indicated that the structure of seminiferous tubules was normal in *Dnah14^-/-^* testis (Fig. 1L). Further Periodic Acid-Schiff (PAS)-hematoxylin staining showed that *Dnah14^-/-^*mice exhibited well-organized seminiferous tubule architecture with normal spermatogenesis progression (Fig. 2A), and the process of sperm head shaping and acrosome formation in *Dnah14^-/-^* mice was similar with wild type mice (Fig. 2B). Thus, disruption of *Dnah14* has little effect on spermatogenesis in mice.

**Figure 2.**
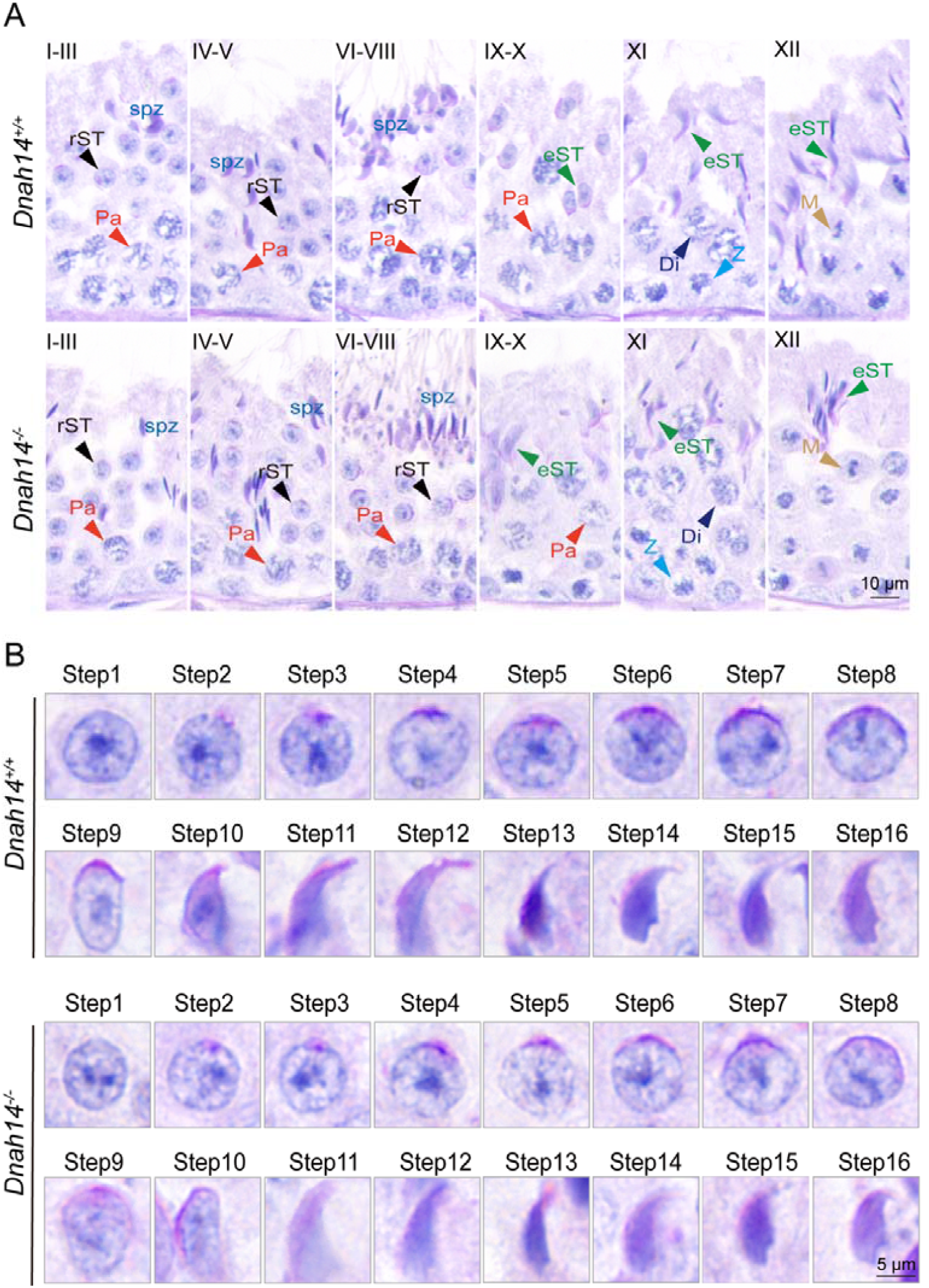
*Dnah14* knockout mice show normal spermatogenesis (A) PAS-hematoxylin staining of *Dnah14^+/+^*and *Dnah14^-/-^* testis sections. Pa: Pachytene spermatocyte, rSt: round spermatid, spz: spermatozoa, M: meiotic spermatocyte, eSt: elongating spermatid, Di: diplotene spermatocytes, Z: zygotene spermatocytes. Scale bars=10 μm. (B) PAS-hematoxylin staining of spermatids at different steps from *Dnah14^+/+^* and *Dnah14^-/-^* mice. Scale bars=5 μm.

Next, we characterized epididymis sections of *Dnah14^-/-^* mice using H&E staining, and found that the sperm count in the *Dnah14^-/-^* caudal epididymis was similar with *Dnah14^+/+^* mice (Fig. 3A-B). Single sperm immunofluorescence staining of key sperm structures showed regular organization of the acrosome (SP56)^32^, mitochondrial sheath (MitoTracker)^31^, fibrous sheath (AKAP4)^33^, and annulus (SEPT4)^31;34^ in *Dnah14^-/-^* spermatozoa (Fig. 3C-F). Further transmission electron microscopy (TEM) analysis confirmed normal “9+2” microtubule structure in *Dnah14^-/-^* sperm flagella (Fig. 3G). Therefore, these results demonstrate that DNAH14 is dispensable for spermatogenesis and sperm structural integrity, indicating that the observed subfertility phenotype likely stems from functional rather than morphological defects in spermatozoa.

**Figure 3.**
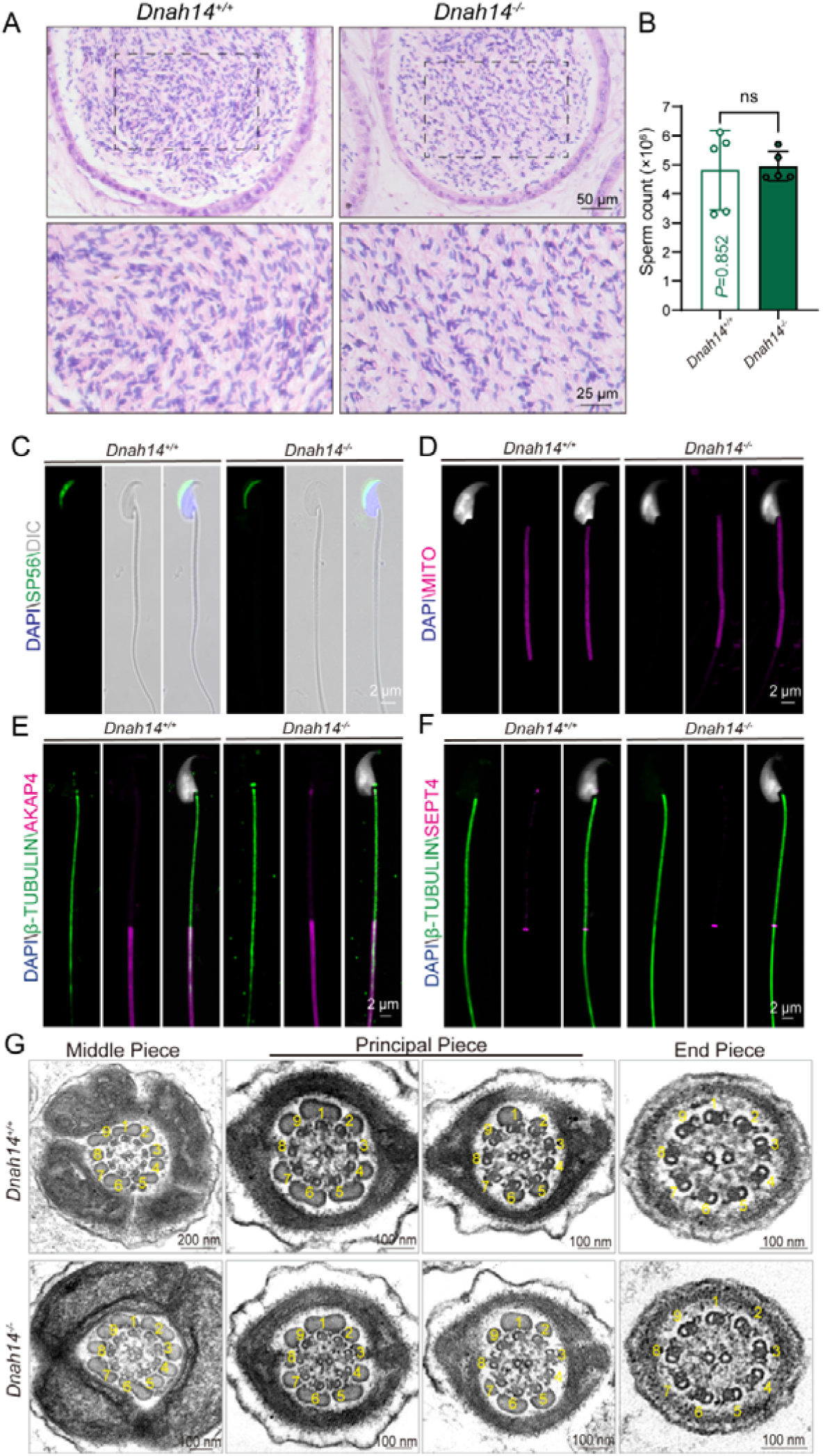
*Dnah14* deletion does not compromise the number and structural integrity of spermatozoa (A) Hematoxylin-eosin (H&E) staining of the cauda epididymis from 2-month-old *Dnah14^+/+^* and *Dnah14^-/-^* male mice. Scale bars=50 μm, 25 μm. (B) Analysis of sperm counts in *Dnah14^+/+^* and *Dnah14^-/-^* male mice (n=5 independent experiments). Mature spermatozoa were extracted from the unilateral cauda epididymis and dispersed in phosphate-buffered saline (PBS). Sperm counts were measured using hemocytometers. Data are presented as the means ± SD. Two-tailed Student’ s t test; ns: no significance. (C) Immunofluorescence staining of acrosomal marker SP56 (green) in sperm from *Dnah14^+/+^* and *Dnah14^-/-^* mice. (D) Immunofluorescence detection of mitochondria (MITO, magenta) in sperm from *Dnah14^+/+^* and *Dnah14^-/-^* mice. (E) Single-sperm immunofluorescence co-staining of fibrous sheath protein AKAP4 (magenta) and TUBULIN (green) in *Dnah14^+/+^* and *Dnah14^-/-^* sperm. (F) Single-sperm immunofluorescence co-staining of annulus protein SEPT4 (magenta) and TUBULIN (green) in *Dnah14^+/+^*and *Dnah14^-/-^* sperm. Scale bars=2 nm (C-F). (G) Ultrastructural analysis of cross-sections of spermatozoa from the cauda epididymis obtained from *Dnah14^+/+^*and *Dnah14^-/-^* mice. Scale bars=200 nm, 100 nm.

### *Dnah14* knockout impairs sperm motility and fertilization capacity

Next, we conducted comprehensive kinematic analyses of sperm motility using computer-assisted sperm analysis (CASA). *Dnah14^-/-^* spermatozoa exhibited significant motility defects compared to controls, including reduced percentages of motile (Fig. 4A) and progressively motile spermatozoa (Fig. 4B). Quantitative analysis revealed velocity parameters were significantly impaired, with decreases in average path velocity (VAP; 111.47 ± 3.54 μm/s vs. 99.06 ± 6.06 μm/s) and straight-line velocity (VSL; 77.77 ± 3.55 μm/s vs. 65.23 ± 5.17 μm/s). Intriguingly, beat-cross frequency (BCF) was elevated (26.79 ± 1.44 Hz vs. 30.57 ± 0.36 Hz), indicating aberrant flagellar dynamics (Fig. 4C-E). Additionally, other parameters, such as linearity (%), curvilinear velocity (μm/s), and straightness (%), showed no significant differences (Fig. 4F-I). To determine whether these motility defects contribute to the subfertility phenotype, we performed IVF assays using *Dnah14^-/-^*spermatozoa. Consistent with the motility impairments, *Dnah14^-/-^*spermatozoa exhibited a significant reduction in fertilization rates (Fig. 4K-L). These results demonstrate that *Dnah14* deficiency specifically disrupts key aspects of sperm motility, including forward velocity parameters and propulsion, ultimately leading to impaired fertilization efficiency and male subfertility.

**Figure 4.**
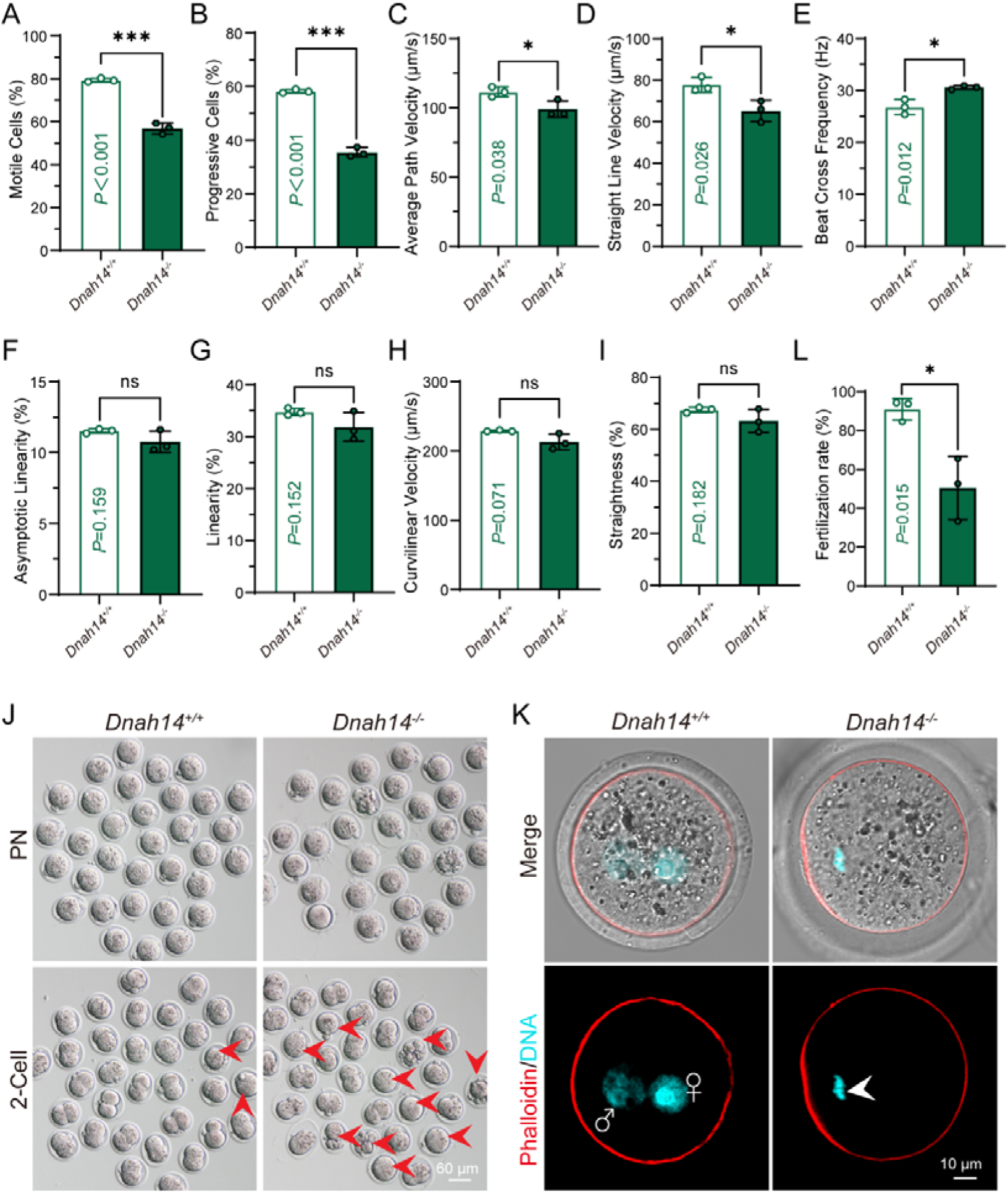
*Dnah14* knockout impairs sperm motility and fertilization capacity (A) The proportion of motile spermatozoa in *Dnah14*^+/+^ and *Dnah14^-/-^*mice (79.34 ± 0.86% vs. 56.91 ± 2.54%, n=3 independent experiments). Data are presented as the means ± SD. Two-tailed Student’s t test; \*\*\**P* < 0.001. (B) The proportion of progressive spermatozoa in *Dnah14*^+/+^ and *Dnah14^-/-^* mice. (58.27 ± 0.62% vs. 35.45 ± 1.77%, n=3 independent experiments). Data are presented as the means ± SD. Two-tailed Student’s t test; \*\*\**P* < 0.001. (C) The average path velocity (VAP) of spermatozoa from *Dnah14*^+/+^ and *Dnah14^-/-^* mice (111.47 ± 3.54 μm/s vs. 99.06 ± 6.05 μm/s, n=3 independent experiments). Data are presented as the means ± SD. Two-tailed Student’s t test; \**P*=0.038. (D) The straight-line velocity (VSL) of spermatozoa from *Dnah14*^+/+^ and *Dnah14^-/-^* mice (77.77 ± 3.55 μm/s vs. 65.23 ± 5.17 μm/s, n=3 independent experiments). Data are presented as the means ± SD. Two-tailed Student’s t test; The beat cross frequency of sperm from *Dnah14*^+/+^ and *Dnah14^-/-^* mice.\**P*=0.026. (E) The beat cross frequency (BCF) of spermatozoa from *Dnah14*^+/+^ and *Dnah14^-/-^* mice (26.79 ± 1.44 Hz vs. 30.57 ± 0.36 Hz, n=3 independent experiments). Data are presented as the means ± SD. Two-tailed Student’s t test; \**P*=0.012. (D) The asymptotic linearity of spermatozoa from *Dnah14*^+/+^ and *DDnah14^-/-^* mice. Data are presented as the means ± SD. Two-tailed Student’s t test; ns: no significance. (E) The straightness of spermatozoa from *Dnah14*^+/+^ and *Dnah14^-/-^* mice. Data are presented as the means ± SD. Two-tailed Student’s t test; ns: no significance. (F) The curvilinear velocity (VCL) of spermatozoa from *Dnah14*^+/+^ and *Dnah14^-/-^* mice. Data are presented as the means ± SD. Two-tailed Student’s t test; ns: no significance. (J) Distribution stages of IVF embryos at pronuclear and two-cell stage from *Dnah14*^+/+^ and *Dnah14^-/-^* groups. 1 × 10^5/mL capacitated sperm were co-incubated with ovulated oocytes in HTF medium for IVF. Arrows indicate abnormal embryos. (K) Representative images of pronuclei stained with Phalloidin (red) and DAPI (cyan) in *Dnah14*^+/+^ and *Dnah14^-/-^* groups after 6 hours post IVF. Arrows indicate chromosomes of unfertilized oocytes. (L) Fertilization rate of *Dnah14*^+/+^ and *Dnah14^-/-^* groups in K. *Dnah14*^+/+^, 90.97% ± 3.19% (n=3 independent experiments, total oocytes=82); *Dnah14^-/-^*, 50.50% ± 9.38% (n=3 independent experiments, total oocytes=85). Data are presented as the means ± SD, \**P*<0.05.

### DNAH14 deficiency reduces sperm flagellar beat amplitude

To investigate the mechanistic basis of reduced sperm velocity in *Dnah14*-deficient males, we performed two-dimensional (2D) analyses of single sperm cells tethered with their heads at the glass surface^35^. Analyses of flagellar beat parameters^36^ showed the characteristic symmetric flagellar beat in *Dnah14^-/-^* spermatozoa (Fig. 5A). Quantitative analysis revealed that *Dnah14^-/-^*sperm exhibited significantly reduced flagellar beat amplitude (Fig. 5B-D), alongside elevated beat frequency (Fig. 5E-F). Additionally, sperm curvature was markedly decreased (Fig. 5H-J), whereas head orientation angle (AngleΘ) remained unaffected (Fig. 5G). These findings mirror observations in *Chlamydomonas*, where inner arm dynein deficiency similarly reduces principal bend amplitude^37^, suggesting evolutionary conservation of dynein-mediated beat regulation. Thus, DNAH14 deficiency impairs sperm motility by reducing sperm flagellar beat amplitude.

**Figure 5.**
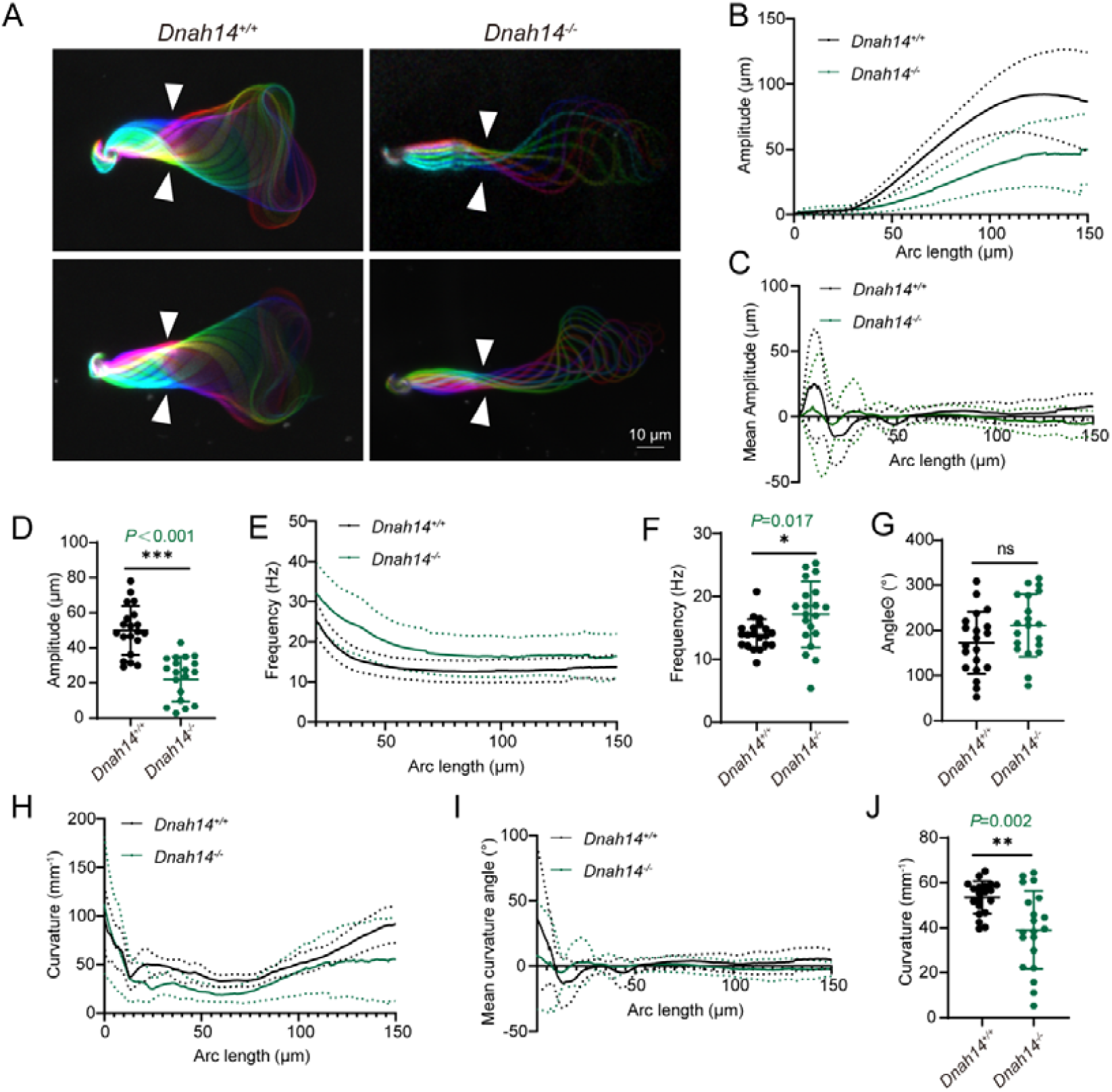
*Dnah14* deficiency reduces sperm flagellar beat amplitude (A) Color-coded time projections of dark-field recordings of head-tethered mouse spermatozoa in two dimensions: The depicted color-coded time span (140 ms) corresponds to one beat cycle of wild-type sperm. Scale bars=10 μm. (B-J) Multiparameter motility analyses of the flagellar beat using the software SpermQ. In all line graphs, solid lines indicate the time-averaged values, and dotted lines the standard deviation for different arc-length positions along the flagellum. Scatter dot plots show the time- and arc-length-averaged values for individual sperm cells (dots) as well as the means values ± SD. (B) Amplitude of *Dnah14*^+/+^ and *Dnah14^-/-^* spermatozoa. Dotted lines indicate the time-averaged values; the standard deviation is presented for different arc-length positions along the flagellum. Data are presented as the means ± SD. (C) Mean flagellar amplitude of *Dnah14*^+/+^ and *Dnah14^-/-^* spermatozoa. Dotted lines indicate the time-averaged values; the standard deviation is presented for different arc-length positions along the flagellum. Data are presented as the means ± SD. (D) Scatter dot plots showing the time- and arc-length-averaged values of the amplitude of the flagellar beat for individual sperm cells (dots). Data are presented as the means ± SD. Two-tailed Student’s t test; \*\*\**P*<0.001. (E) Frequency of *Dnah14*^+/+^ and *Dnah14^-/-^* spermatozoa. Dotted lines indicate the time-averaged values; the standard deviation is presented for different arc-length positions along the flagellum. Data are presented as the means ± SD. (F) Scatter dot plots showing the time- and arc-length-averaged values of the frequency of the flagellar beat for individual sperm cells (dots). Data are presented as the mean ± SD. Two-tailed Student’s t test; \**P*=0.017. (G) Head angle in space (Θ) of flagellar beat in *Dnah14*^+/+^ and *Dnah14^-/-^* spermatozoa. Data are presented as the means ± SD. Two-tailed Student’s t test; ns: no significance. (H) Flagellar curvature from *Dnah14*^+/+^ and *Dnah14^-/-^* spermatozoa. Dotted lines indicate the time-averaged values, the standard deviation is presented for different arc-length positions along the flagellum. Data are presented as the means ± SD. (I) Mean curvature angles of *Dnah14*^+/+^ and *Dnah14^-/-^* spermatozoa. Dotted lines indicate the time-averaged values; the standard deviation is presented for different arc-length positions along the flagellum. Data are presented as the means ± SD. (J) Scatter plot shows the time- and arc-length-averaged values of curvature angle for individual sperm cells (dots). Data are presented as the means ± SD. Two-tailed Student’s t test; \*\**P*=0.002.

### Dose-dependent rescue of fertilization capacity in DNAH14-deficient male mice

To investigate whether DNAH14 deficiency affects the integrity of other dynein arm components, we performed immunofluorescence analysis of two critical dynein subunits: DNAH6 (an inner dynein arm protein) and DNAH17 (an outer dynein arm protein)^38^. Both proteins maintained normal subcellular localization patterns and expression levels in *Dnah14^-/-^* spermatozoa (Fig. 6A-B), suggesting that DNAH14 loss specifically impairs sperm movement without causing global disruption of axonemal dynein arms.

**Figure 6.**
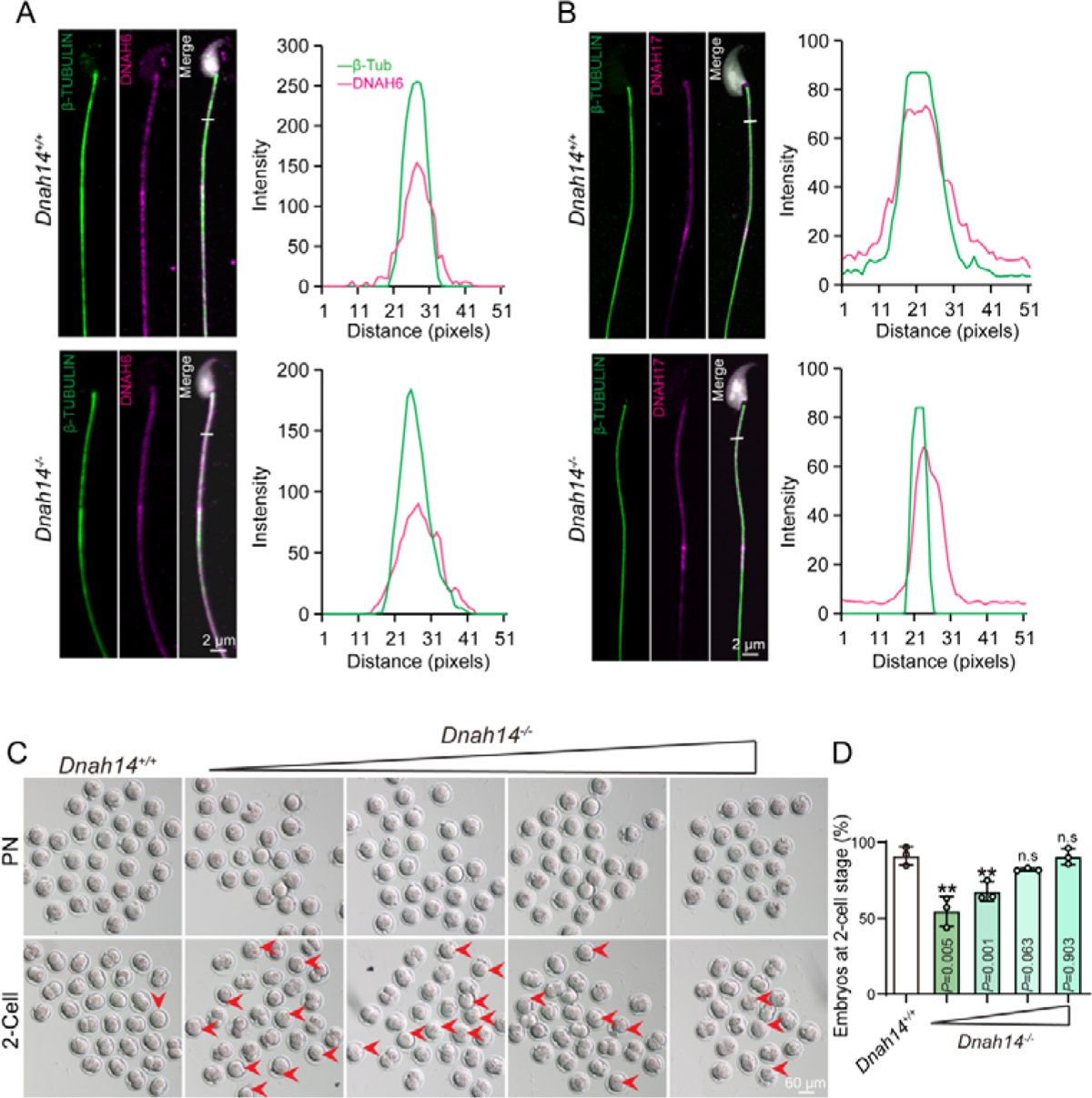
Dosage-dependent rescue of fertilization capacity in DNAH14-deficient male mice (A) Single-sperm immunofluorescence co-staining of annulus protein DNAH6 (magenta) and TUBULIN (green) in *Dnah14^+/+^* and *Dnah14^-/-^* spermatozoa. Line-scan analysis (white line) was performed using ImageJ software. (B) Single-sperm immunofluorescence co-staining of annulus protein DNAH17 (magenta) and TUBULIN (green) in *Dnah14^+/+^* and *Dnah14^-/-^* spermatozoa. Line-scan analysis (white line) was performed using ImageJ software. (C) Developmental progression of IVF embryos at the pronuclear and two-cell stages in *Dnah14*^+/+^ and *Dnah14^-/-^* groups. For in vitro fertilization, ovulated oocytes were co-incubated with 1 × 10^5^/mL capacitated *Dnah14*^+/+^ sperm, or 1-4 × 10^5/mL capacitated *Dnah14^-/-^* sperm in HTF medium. Arrows indicate abnormal embryos. (D) Percentage of embryos at the two-cell stage from *Dnah14*^+/+^ and *Dnah14^-/-^* groups. *Dnah14*^+/+^, 91.17% ± 3.52% (n=3 independent experiments, total oocytes=72); *Dnah14^-/-^* (from left to right), 54.63% ± 5.68% (total oocytes=63), 67.63% ± 3.69% (total oocytes=70), 82.10% ± 0.46% (total oocytes=67), 90.57% ± 2.99% (total oocytes=66), n=3 independent experiments. Data are presented as the means ± SD, ** *P*<0.01, ns: no significance.

To examine whether the observed biomechanical defects could be functionally compensated, we conducted IVF assays using progressively increasing concentrations of *Dnah14^-/-^* spermatozoa. Notably, while control sperm achieved maximal fertilization at standard concentrations, *Dnah14^-/-^* spermatozoa required significantly higher concentrations (Fig. 6C) to restore fertilization rates to near-normal levels (Fig. 6D). This dose-dependent rescue suggests that quantitative enhancement of sperm-egg encounters can partially overcome the qualitative motility impairments caused by DNAH14 deficiency.

## Discussion

Asthenozoospermia constitutes a predominant etiological factor in male infertility^39^. While the pathogenesis of this condition is recognized as multifactorial, encompassing genetic, epigenetic, and environmental determinants, the underlying mechanisms remain idiopathic in a substantial proportion of clinical presentations^40;41^. Here, we establish DNAH14 as a novel genetic contributor to asthenozoospermia through its essential role in regulating sperm flagellar motility. The generation and characterization of *Dnah14* knockout mice revealed a distinct asthenozoospermic phenotype (Fig. 4A-D) that differs fundamentally from previously reported dynein-related sperm defects^42;43^. While mutations in other DNAH family members typically cause either complete sperm immotility or MMAF^11;18^, DNAH14 deficiency results in a unique kinematic profile characterized by significantly reduced flagellar beat amplitude coupled with increased beat frequency (Fig. 5B-F), and the knockout of *Dnah14* has no effect on sperm morphology and axonemal ultrastructure (Fig 3G). This finding suggests the existence of a previously unrecognized subclass of asthenozoospermia in clinical populations, where patients may present with normal semen parameters but specific impairments in progressive motility due to altered flagellar dynamics.

The preservation of other dynein arms (DNAH6 and DNAH17) in *Dnah14* knockout spermatozoa indicates that DNAH14 operates within a distinct functional module of the axonemal complex, primarily influencing power stroke generation (Fig. 6A-B). DNAH14 may function as a specialized regulator of flagellar biomechanics. This specificity likely explains why DNAH14 deficiency produces a milder phenotype compared to mutations affecting more fundamental components of the dynein regulatory system^17;20^. The evolutionary conservation of this mechanism, as evidenced by similar findings in *Chlamydomonas*^37;44^, underscores the fundamental importance of inner arm dyneins in controlling flagellar bending patterns.

Although *Dnah14* knockout cause male subfertility (Fig. 1G), increased sperm concentration can partially rescue the fertilization deficit in DNAH14-deficient males (Fig. 6C-D) suggests practical therapeutic approaches for analogous human cases. For men with similar genetic defects, relatively simple interventions such as intrauterine insemination with concentrated sperm preparations could represent an effective first-line treatment, potentially avoiding the need for more invasive and expensive intracytoplasmic sperm injection (ICSI) procedures. This approach would be especially valuable in resource-limited settings where advanced assisted reproduction technologies are less accessible. Thus, DNAH14 is identified as a specialized regulator of sperm flagellar biomechanics, and our finding provides new insights into the genetic basis of male infertility.

## Supporting information

Trachea cilia and lung cilia appear normal in Dnah14-/- mice

## Author contributions

Y.L., Y.Z, Y.Q performed experiments and data collection. J.Z., H.X., B.W. carried out computation and statistical analysis. Y.L., L.W., W.L., C.L. wrote the manuscription with contributions from all authors. L.W., W.L., C.L., X.T. supervised the project. All authors reviewed and approved the final manuscript.

## Acknowledgements

We would like to express our sincere gratitude to all those who have supported and guided this research.

