## Supplementary figures and images for "DNAH14 deficiency impairs sperm motility by reducing flagellar beat amplitude"

### Trachea cilia and lung cilia appear normal in Dnah14-/- mice

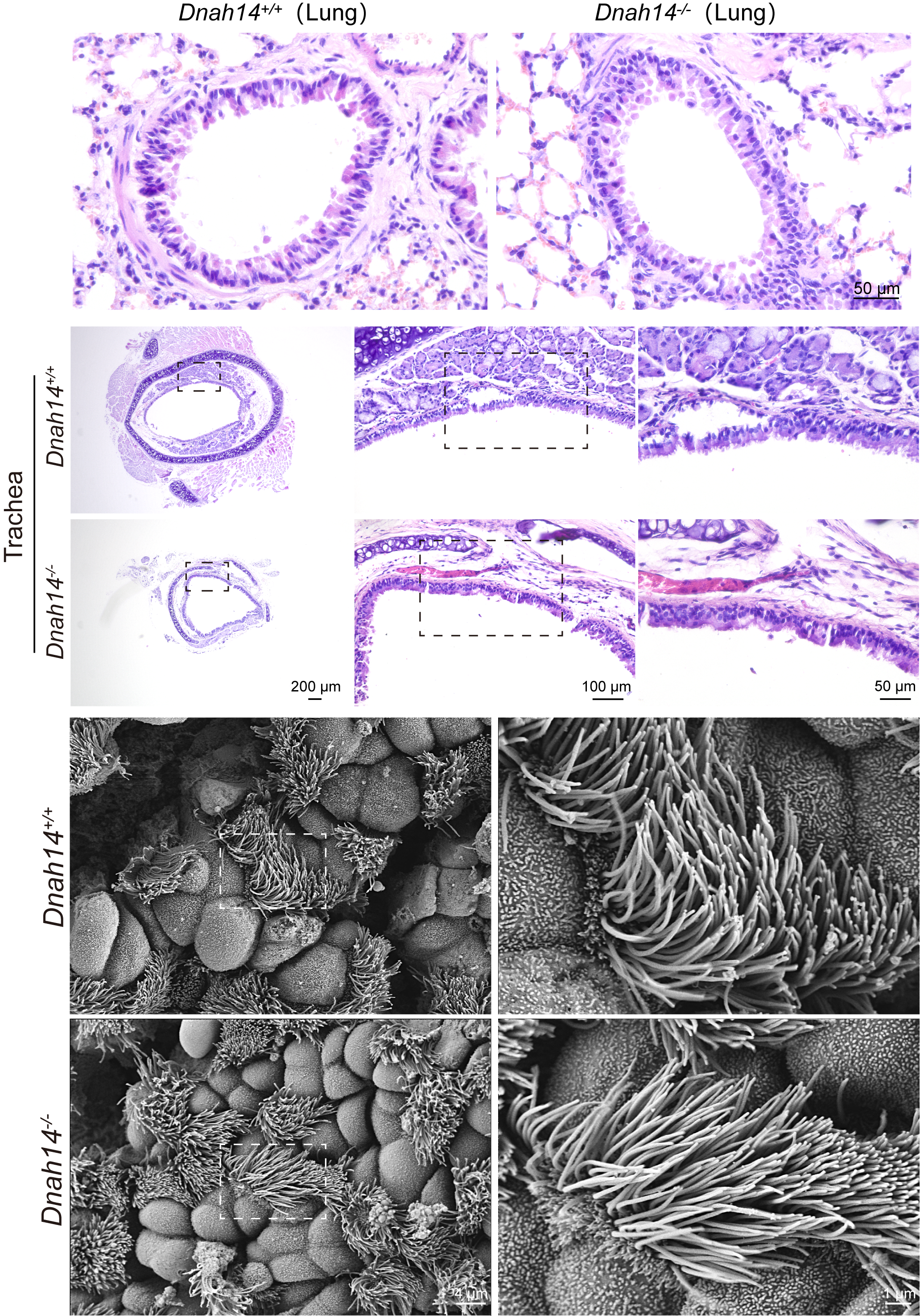
